# Reassessing the association of VDR and its polymorphisms with tuberculosis in global populations

**DOI:** 10.1101/2023.12.09.570914

**Authors:** Debashruti Das, Gyaneshwer Chaubey

## Abstract

**Background:** Vitamin D is a hormone that regulates the calcium homeostasis of the body. Besides this classical function, it is also regarded as an important immunomodulator. Most active Vitamin D actions are mediated through the Vitamin D receptor (VDR), a transcription factor and also a member of the nuclear receptor superfamily. In this study, we explored the phylogeographic attributes of the four most well-known polymorphisms of the VDR gene namely rs7975232 (ApaI), rs731235 (TaqI), rs1544410 (BsmI), rs2228570 (FokI) and also evaluated their association with the incidence of tuberculosis in global populations. This study integrated several in-silico approaches on population databases to evaluate the pattern of distribution, linkage and selection patterns of these SNPs.

**Results:** The ancestral alleles of rs7975232, rs731235, and rs1544410 are still present in over 50% frequency in modern human populations. These SNPs also have a very strong linkage disequilibrium among themselves in all population groups but no haplotype blocks are seen in South Asian populations constituting these polymorphisms. The selection results reveal a negative Tajima’s D value in West and East Eurasian populations suggesting positive selection in these regions… In correlation studies, we found no association between the incidence of tuberculosis and the allele or genotype frequency of these four SNPs.

**Conclusion:** The four SNPs of VDR behave differently in South Asian populations as compared to West and East Eurasian populations but no significant association was found with the incidence of tuberculosis in global populations.

## Introduction

Pathogenic diseases are the second leading cause of death worldwide after cardiovascular diseases, with Tuberculosis (TB), being the number one cause on a global scale and have been among the top-ranked dangerous diseases by WHO over decades. The disease is caused by the members of the Mycobacterium tuberculosis complex, including *Mycobacterium tuberculosis*, the agent of TB in humans. Decoding the 4Mbp genome of *M. tuberculosis* helped in the reconstruction of the history of the pathogen. ^[3]^ It emerged as a potent human pathogen about 70,000 years ago in Africa and has spread worldwide with human migration following an “evolutionary bottleneck” in the disease. ^[5]^ Though it typically attacks the lungs, the bacteria also affect the brain, spine and kidneys. One-third of the worldwide population is suspected to be affected by the bacteria ^[8]^. Still, only 10% of those develop active TB ^[1,2,4,6],^ and even then, the symptoms (cough, fever, cold sweats, weight loss etc.) may be mild for many months leading to delay in seeking treatment.

Despite the exhaustive unifying effort, TB still accounts for a huge burden of mortality and morbidity globally. Though TB is present worldwide, developing countries account for a disproportionate share of the disease burden, especially India, Indonesia, China, Nigeria, Pakistan, and South Africa and remain a significant cause of death, particularly in immuno-compromised individuals like those affected with AIDS. ^[22]^ (Tuberculosis accounted for 35% of mortality in HIV-infected people in 2015.)

In order to manage any disease, it is of utmost importance to understand its pathogenesis and progression along with clinical manifestation. Tuberculosis spreads from person to person through airborne aerosol droplets. Droplets smaller than 5um reach the distal airways, phagocytosed by immune cells like macrophages, dendritic cells, and neutrophils. Evidence of Mtb-infected non-myelocytic cells like type II alveolar epithelial cells has also been found. (Bussi and Gutierrez, 2019; Teitelbaum et al. 1999) The bacilli in larger droplets that get stuck in the upper passageways invade a type of specialised epithelial cells called Microfold cells (M cells). (Bussi and Gutierrez, 2019; Teitelbaum et al. 1999) These cells, in turn, elicit an inflammatory response locally that induces the formation of a multicellular aggregate--- granuloma, the hallmark of tuberculosis and a very complex environment for the growth of the bacilli.

**Fig 1:**
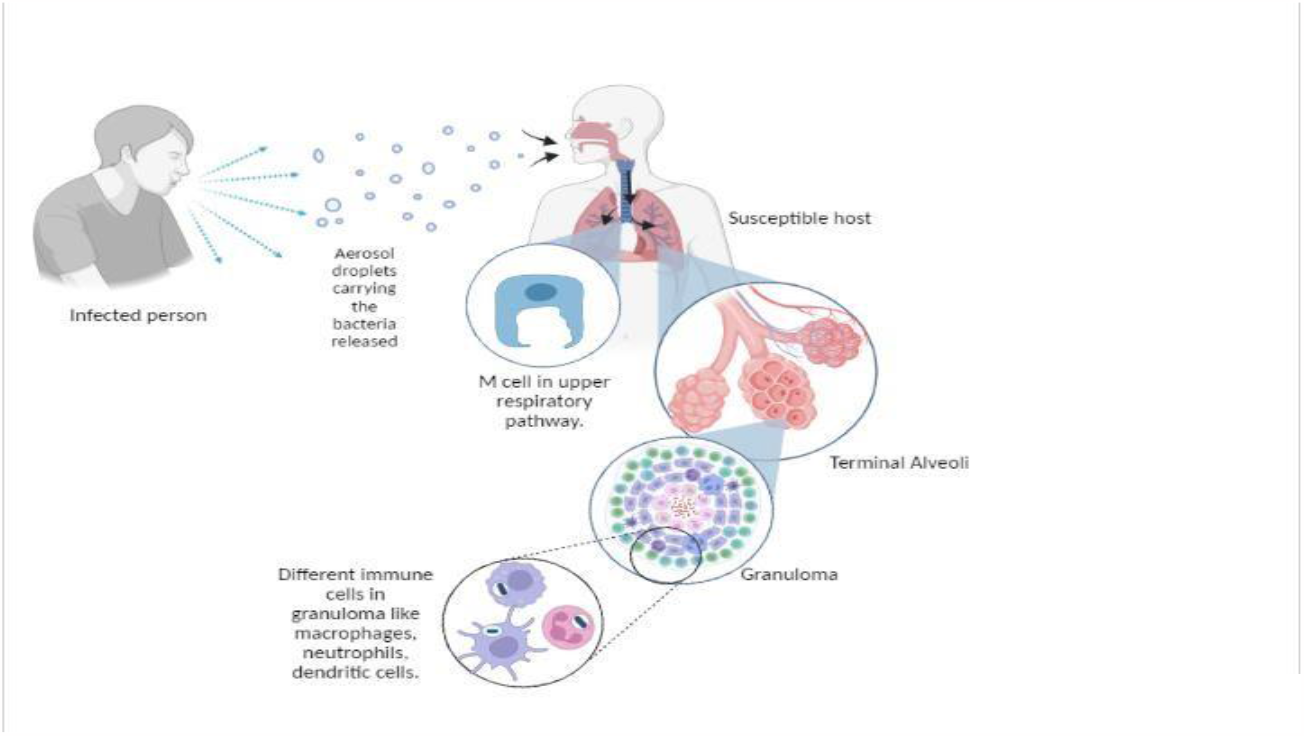
Pathogenesis of Tuberculosis. The pathogen ridden aerosol droplets disseminated from the infected person reaches a susceptible host and reaches the lungs. Some pathogens are managed by the M cells, modified epithelial cells in the upper respiratory tract. When the pathogen reaches terminal alveoli, it elicits an immune response and a granuloma is formed which contains a plethora of immune cells.

It is very common to target the receptor(s) responsible for pathogen entry in case of infectious disease to control the development of disease and find alternative methods of treatment. In this case, this approach seems ambiguous as there is a plethora of immune and non-myelocytic cells involved which again harbours multiple receptors for pathogen entry. Hence, we studied a common factor which somehow affects the efficiency of these immune cells. Many previous studies have reported the role of Vitamin D as an immune modulator besides being involved in calcium homeostasis. Also, high TB burdens countries like Indonesia, India, Kenya, and others have repeatedly reported low levels of serum Vit D. ^[11]^ It has been reported that a predominant percentage of Vit D deficient people not only suffer from rickets ^[27-29]^ or osteoporosis ^[30-31]^ but also cancer^[32,33]^, cardiovascular diseases^[34,35]^, diabetes^[36,37]^, infectious diseases like tuberculosis^[10,11,13,15]^ and even Covid 19.^[7]^ Most immune cells, like macrophages and monocytes, have Vitamin D receptors on their cell surface. Vitamin D is a steroid hormone that is known to regulate a multitude of functions in the human body. Most importantly, it is involved in the mechanisms of innate and acquired I immunity and produces antimicrobial agents like cathelicidin and humanβ-defensin-2. It also regulates the expression of genes that is responsible for the intracellular destruction of pathogens.^[7]^ The biologically active form of Vit D is a double hydroxylated compound 1,25(OH)_2_D, also called calcitriol. This binds to vitamin D binding protein, which in turn expresses its function in various target tissues through the nuclear Vitamin D receptor (VDR). The gene translating into VDR is present on chromosome 12q12-q14. VDR aids in the transcription of genes that help to eliminate the bacteria within the macrophages. The VDR gene regulates innate immunity by modulating macrophage and monocyte activity. ^[6,8,10,11]^ Vit D helps to increase the fusion of phagosome and lysosome in macrophages infected with the bacteria.^[9,10,11]^

**Fig 2:**
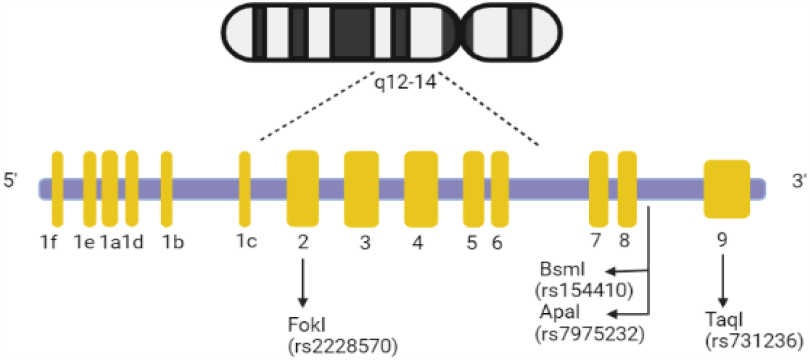
VDR gene in genomic location. Structure of VDR gene showing the position of the polymorphisms studied here.

Hence, decreased levels of Vit D and/or impaired receptor structure and function might lead to compromised immunity towards the pathogen. [8,10] Genetic polymorphisms in the VDR gene might affect the structure of the VDR protein or cause defects in its regulation or function, which in turn can alter the biological functions of Vitamin D. There are more than 25 VDR polymorphisms in the 5’ regulatory region, 3’UTR and other coding regions[18], four being the most relevant among them namely ApaI(intron VIII), BsmI (intron VIII), TaqI (Exon IX) and FokI (Exon II). ^[17,18],^ which will be further analysed in the following sections.

## Methods

The allele frequency of the variants of VDR gene was calculated using the NGS data from Pagani et.al. ^[38]^ and was validated using the 1000 Genome data. The spatial maps depicting the allele distribution for ancient and modern populations were generated using the PGG toolkit^[45]^, a repository of several modern and ancient human population databases (including 1000 genome) covering over 200 million SNVs.For further phylogeographic analysis, the various populations from the NGS data were grouped into West Eurasia, East Eurasia and South Asia for easy comparison using PLINK 1.9.^[39]^ East Eurasia contained populations of South East Asia and Siberia; West Eurasia had Europeans, Caucasus and populations from Central Asia and West Asia; while South Asia had samples from South Asia Mainland and Island. Linkage Disequilibrium maps were constructed through Haploview.^[40]^ and LDLink NIH. For each of the three population groups, the PLINK files were converted into fasta format (.ped to IUPAC) using a PERL script. The unphased output fasta files were used as the input files for DNASP v6 for phasing and generating input files for Arlequin and Network. Pairwise genetic differences within and between populations, Nei’s genetic distance were calculated in Arlequin 3.5^[42]^ and plotted using R v 3.1.^[43]^ These results were validated by creating a phylogenetic tree using Mega X. Network v5 and Network Publisher^[44]^ were used to construct median-joining haplotype network to analyze the haplotype distribution and diversity in different population groups.

In order to correlate the incidence of tuberculosis with the polymorphisms in the VDR gene, number of tuberculosis cases for the year 2020 was extracted from WHO database for each country. The number of cases were then grouped and the average was calculated for each population group. These average values were then plotted against the minor allele frequencies as well as genotype frequencies of each SNPs to explore their association.

## Results

### Comparison of the SNPs between ancient and modern human populations

The ancestral allele for ApaI, BsmI, TaqI and FokI is C, C, A and A, respectively. The ancestral allele of 3 out of 4 SNPs, namely ApaI (rs7975232), BsmI (rs1544410) and TaqI (rs731236) still holds a frequency of approximately 50% and above in modern global populations with highest proportion in East Asian populations. The trend is just the opposite in case of FokI (rs2228570) where the modern populations show very little to no presence of the ancestral allele.

**Fig 3A:**
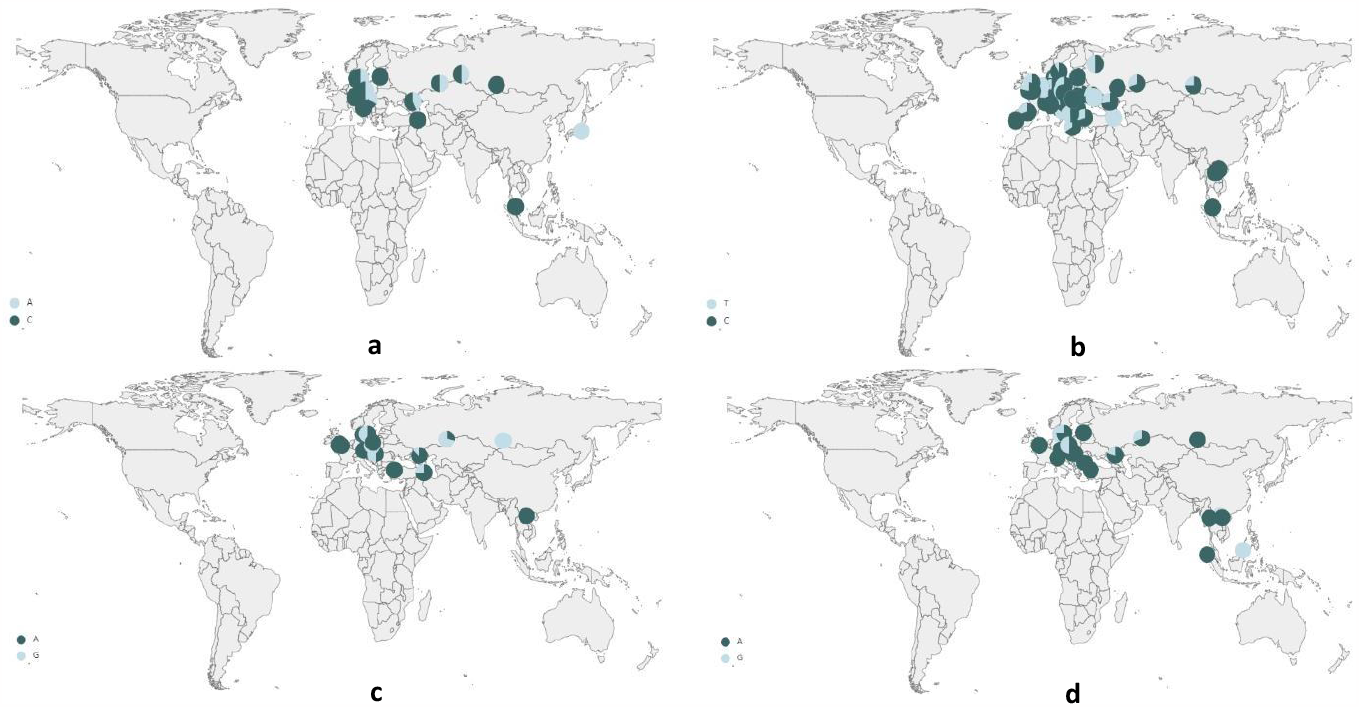
Allele distribution in ancient human genomes (a) ApaI, (b) BsmI, (c)TaqI, (d) FokI

**Fig 3B:**
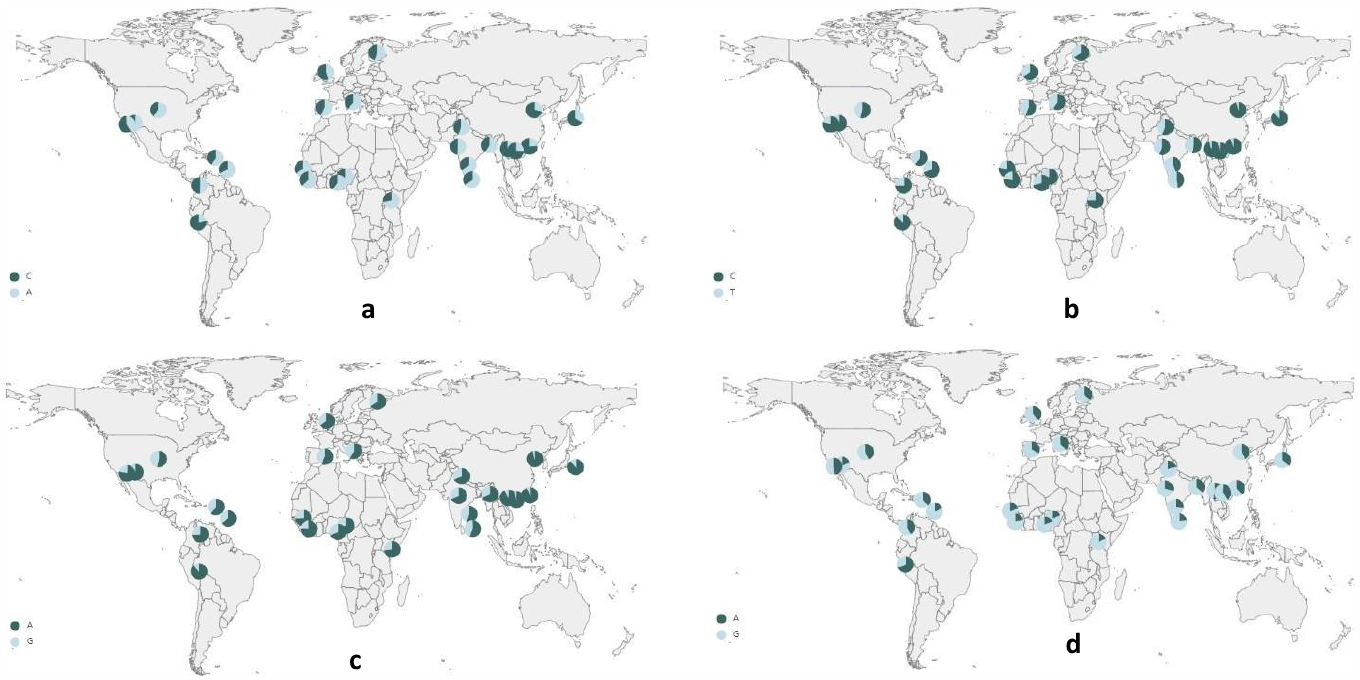
Allele distribution in modern human genomes (a) ApaI, (b) BsmI, (c)TaqI, (d) FokI

### Linkage disequilibrium

The degree of linkage disequilibrium varies significantly within and among loci and populations. This software displays the LD map as per default settings based on D’ and LOD score values while the values inside the boxes are the r^2^ values. The white boxes mean D’<1 and LOD<2. The shades of pink or red indicate a D’<1 and LOD score of ≥ 2 whereas bright red means a D’ value of 1 and LOD score ≥ 2.

In these LD maps, we see blocks are formed, which represent regions (shown in straight lines) that are inherited without much recombination and punctuated by regions (shown in dashed lines) where there are historical traces of recombination between SNPs.

As we can see in Fig. 4. all three population groups show ApaI, TaqI and BsmI in high LD with each other, while FokI is not in high linkage disequilibrium with any of the SNPs in the 34kb span. The LD map of the West and East Eurasian population show that ApaI, TaqI and BsmI are included in haplotype blocks of high LD (11 kb and 14 kb block respectively) and are interrupted by adjacent regions on both sides showing possible recombination. The same analysis was carried out with 1000 genome data and the results (Supplementary data LD 1000 genome compiled.xsv) were similar to this analysis. From Fig. 5, it is seen clearly that the three SNPs contribute to six and seven haplotypes in West and East Eurasian populations, respectively. For both these population groups, the A/T allele of TaqI is predominantly present in the major haplotypes (0.833 in West Eurasian population and 0.857 in East Eurasian populations). For ApaI, the ‘A/T’ allele is slightly more prevalent than ‘C/G’ allele (0.666 vs 0.333 in West Eurasian populations and 0.571 vs 0.428 in East Eurasian populations) whereas for BsmI the scenario is quite the opposite. The C/G allele is major in the haplotypes for both the population groups. (0.666 in West Eurasians and 0.857 in East Eurasians). In South Asia, such haplotype blocks do not include any of the four SNPs suggesting that the South Asian population must have a higher degree of recombination and variability for these SNPs.

**Fig 4:**
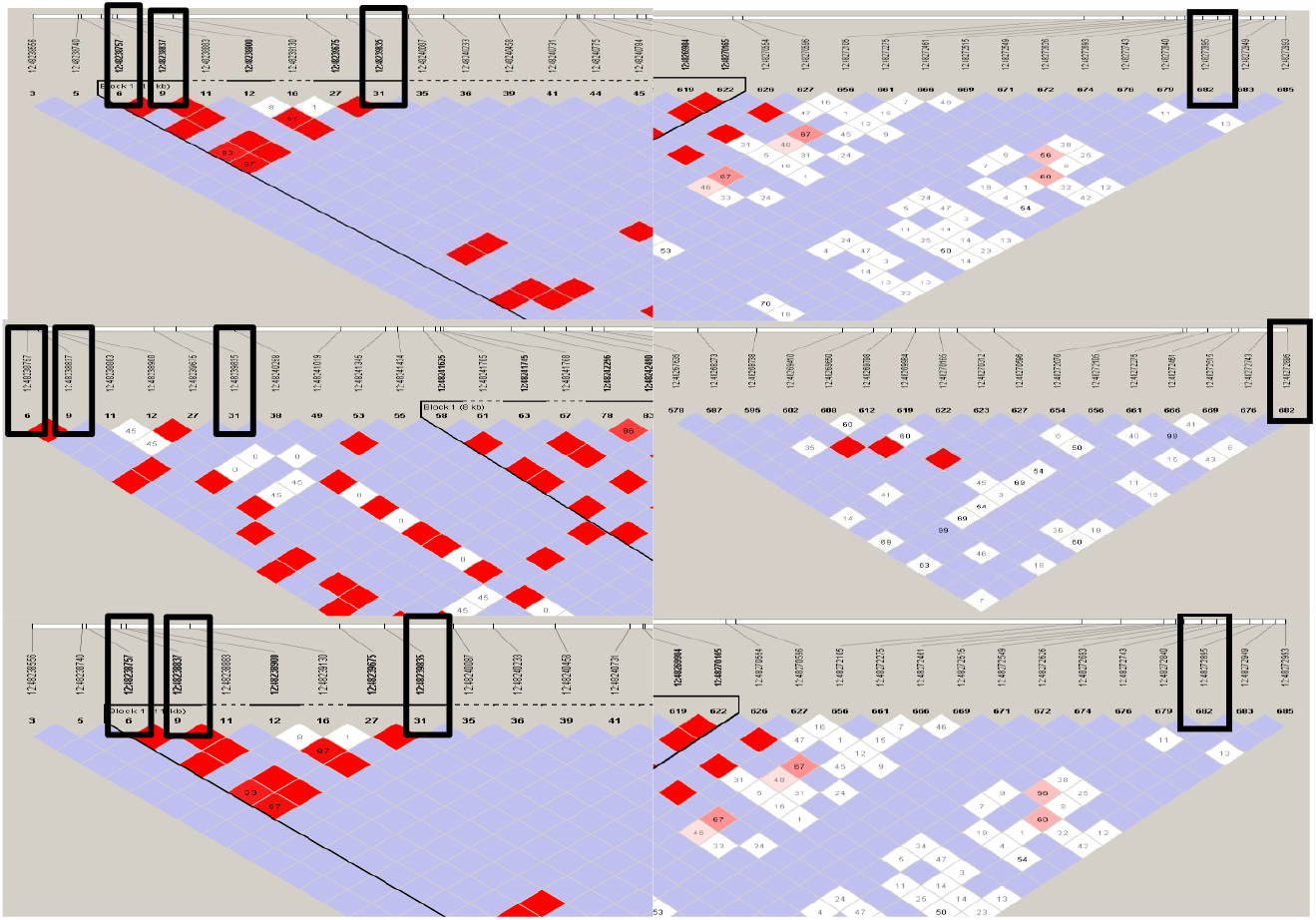
Linkage disequilibrium map of the four VDR polymorphisms in (a) West Eurasian (b) South Asian and (c) East Eurasian populations

**Fig 5:**
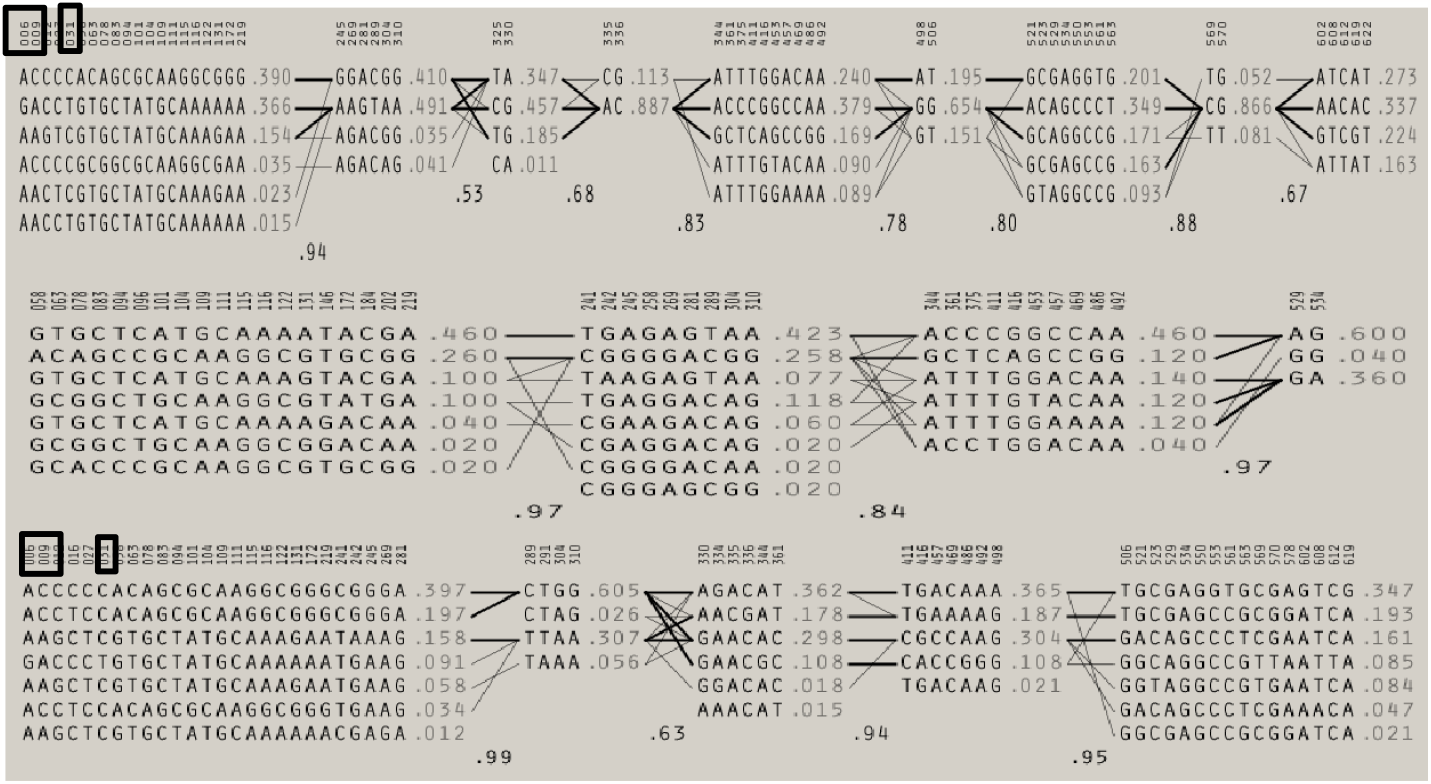
Haplotype diversity of the four SNPs in (a) West Eurasian (b) South Asian (c) East Eurasian populations

**Table 1:** Table showing linkage disequilibrium values of the four SNPs in different populations.

|  | East Eurasia |  |  |  | South Asia |  |  |  | West Eurasia |  |  |  |
| --- | --- | --- | --- | --- | --- | --- | --- | --- | --- | --- | --- | --- |
|  | D' | LOD | R<sup>2</sup> | Confidence interval | D' | LOD | R<sup>2</sup> | Confidence interval | D' | LOD | R<sup>2</sup> | Confidence interval |
| rs731236<br>rs 7975232 | 1.0 | 9.53 | 0.182 | 0.81-1.0 | 1.0 | 3.5 | 0.48 | 0.62-1.0 | 1.0 | 28.61 | 0.44 | 0.93-1.0 |
| Rs731236<br>Rs1544410 | 1.0 | 31.11 | 0.82 | 0.91-1.0 | 1.0 | 6.39 | 0.72 | 0.77-1.0 | 0.97 | 64.31 | 0.91 | 0.93-1.00 |
| Rs731236<br>Rs2228570 | 0.028 | 0.0 | 0.0 | 0.0-0.5 | 0.27 | 0.09 | 0.01 | 0.02-0.76 | 0.02 | 0.01 | 0.0 | -0.01-0.26 |
| Rs7975232<br>Rs1544410 | 1.0 | 11.62 | 0.22 | 0.85-1.0 | 1.0 | 5.27 | 0.66 | 0.74-1.0 | 1.0 | 29.23 | 0.46 | 0.93-1.0 |
| Rs7975232<br>Rs2228570 | 0.024 | 0.01 | 0.001 | -0.01-0.18 | 0.31 | 0.22 | 0.038 | 0.03-0.71 | 0.086 | 0.2 | 0.005 | 0.0-0.25 |
| Rs1544410<br>Rs2228570 | 0.04 | 0.01 | 0.0 | 0.0-0.34 | 0.25 | 0.07 | 0.017 | 0.02-0.76 | 0.011 | 0.0 | 0.0 | -0.01-0.25 |

### Median-Joining (MJ) Haplotype Network Analysis

The network diagram constructed using the median-joining algorithm produced four major haplotype groups. The two haplotypes on the top corner show a predominance of East Eurasian samples, while the bottom two show a West Eurasian predominance. Three out of four SNPs under study i.e ApaI, BsmI and TaqI are seen to be responsible for the rise of West Eurasian-specific haplotypes. The other SNP, FokI, shows the highest variability and gave rise to a number of singletons giving rise to an extremely complex network (Supplementary Fig 1.). Hence, it has been eliminated from the present network analysis. South Asia attributed very less to the major haplotype groups as was predicted from the linkage disequilibrium analysis, while Siberians contributed to most of the haplotype groups.

**Fig 6:**
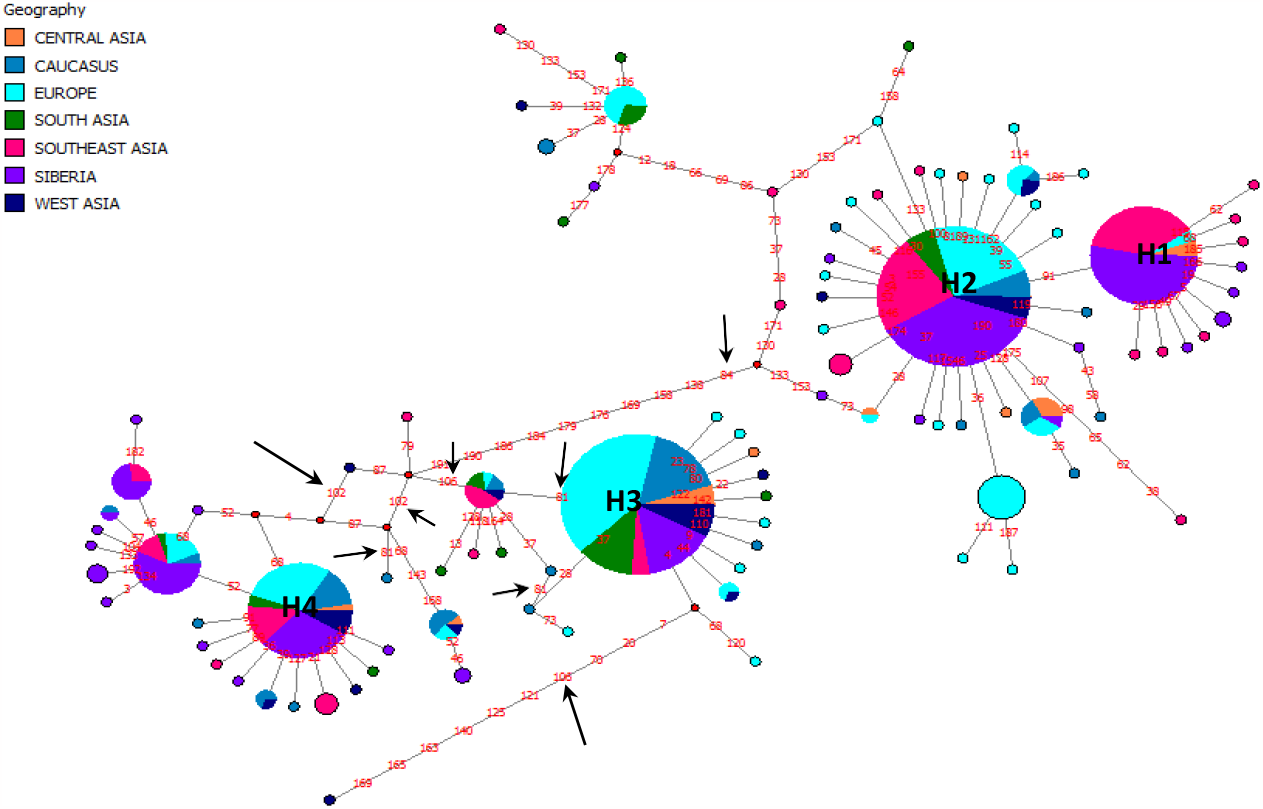
Network Analysis using the median joining algorithm including ApaI, BsmI and TaqI polymorphisms of the VDR gene.

### Pairwise differences in populations

To understand the differences between population groups for this SNP, a pairwise differences matrix was made. The matrix is divided diagonally by various hues of orange squares which represent the within population pairwise differences for any polymorphism. The upper half of the matrix signifies the between population pairwise differences illustrated with different gradients of green. Nei’s distance can be studies from the lower half of the matrix represented by different shades of blue. The bright or dull hues of different colours coincide with the degree of pairwise differences referenced by a legend provided alongside. The pairwise difference within population based on stepwise mutation method was highest in South Asia suggested by the deep orange colour block and least in Southeast Asia represented by a white square for all the four SNPs. Almost similar patterns of pairwise differences between populations are seen for ApaI, TaqI and BsmI polymorphisms. For these SNPs highest differences are seen between Southeast Asians and Caucasians at shown by the deep green colour (values are ∼10.5 for ApaI and TaqI while ∼12.2 for BsmI) followed by South Asia and Southeast Asia. The same is reflected in the Nei’s distance. South Asian and Western Eurasian populations show lower pairwise differences for this SNPs. For FokI, highest pairwise differences were noticed between South Asians and Europeans (value ∼5.45) followed by Southeast Asians and different West Eurasian populations which is also reflected in Nei’s genetic distance. Within population heterogeneity is also high for this SNP for all populations except for Southeast Asians and Siberians.

The neighbour joining phylogenetic tree (Supplementary Fig.2) that was created validated these results. South Asians were found to be closer to West Eurasian populations especially the Caucasians while the East Eurasian Group was placed further away.

**Fig 7:**
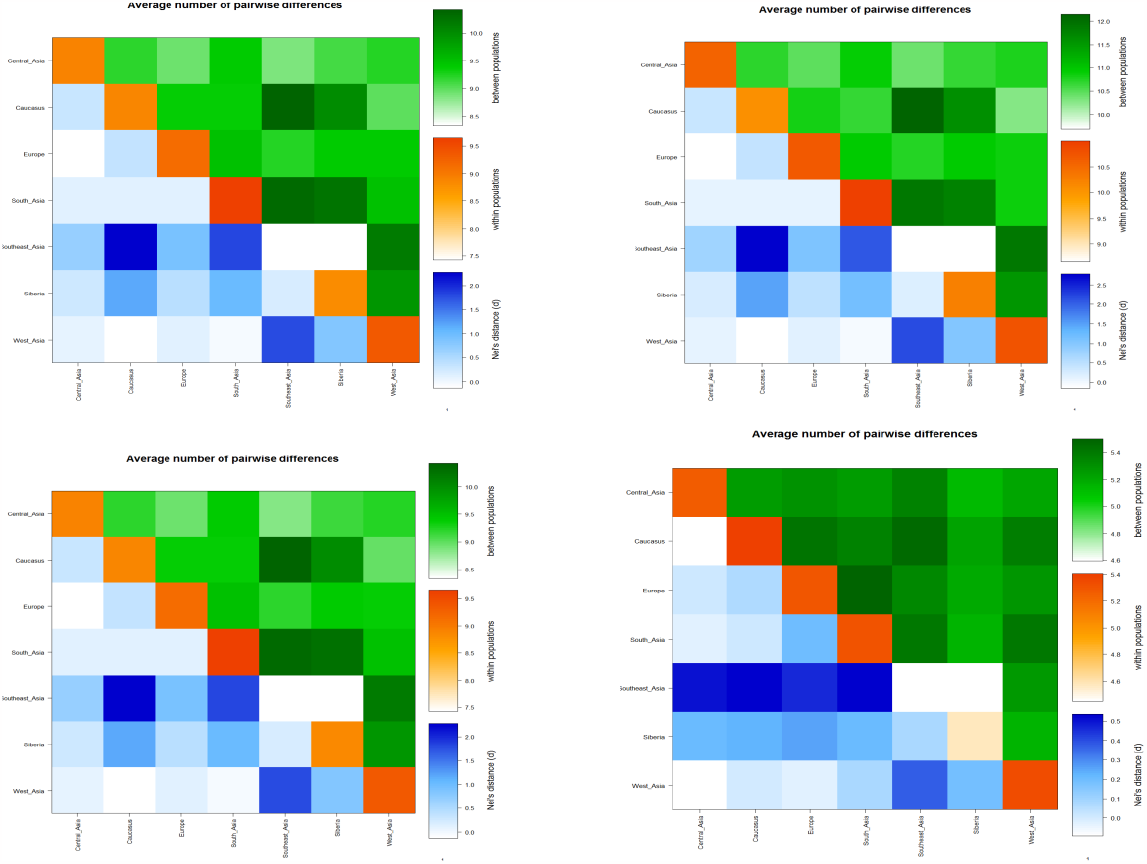
Pairwise difference matrix within population (diagonal orange blocks), between populations (green blocks) and Nei’s distance between populations with respect to (a) ApaI, (b) BsmI, (c) TaqI and (d) FokI.

### Correlation analysis

A correlation analysis was carried out to understand the association between the allele (Fig 8a) or genotype frequency (Fig 8b) and average number of tuberculosis cases in global population. Our results showed no particular association between the two and the null hypothesis was accepted. South Asians had a high number of tuberculosis cases and America, Central Asia, Europe reported lower tuberculosis cases despite low or high allele frequencies. This result points out that the above polymorphisms have no direct effect on tuberculosis susceptibility singly.

**Fig 8(a).**
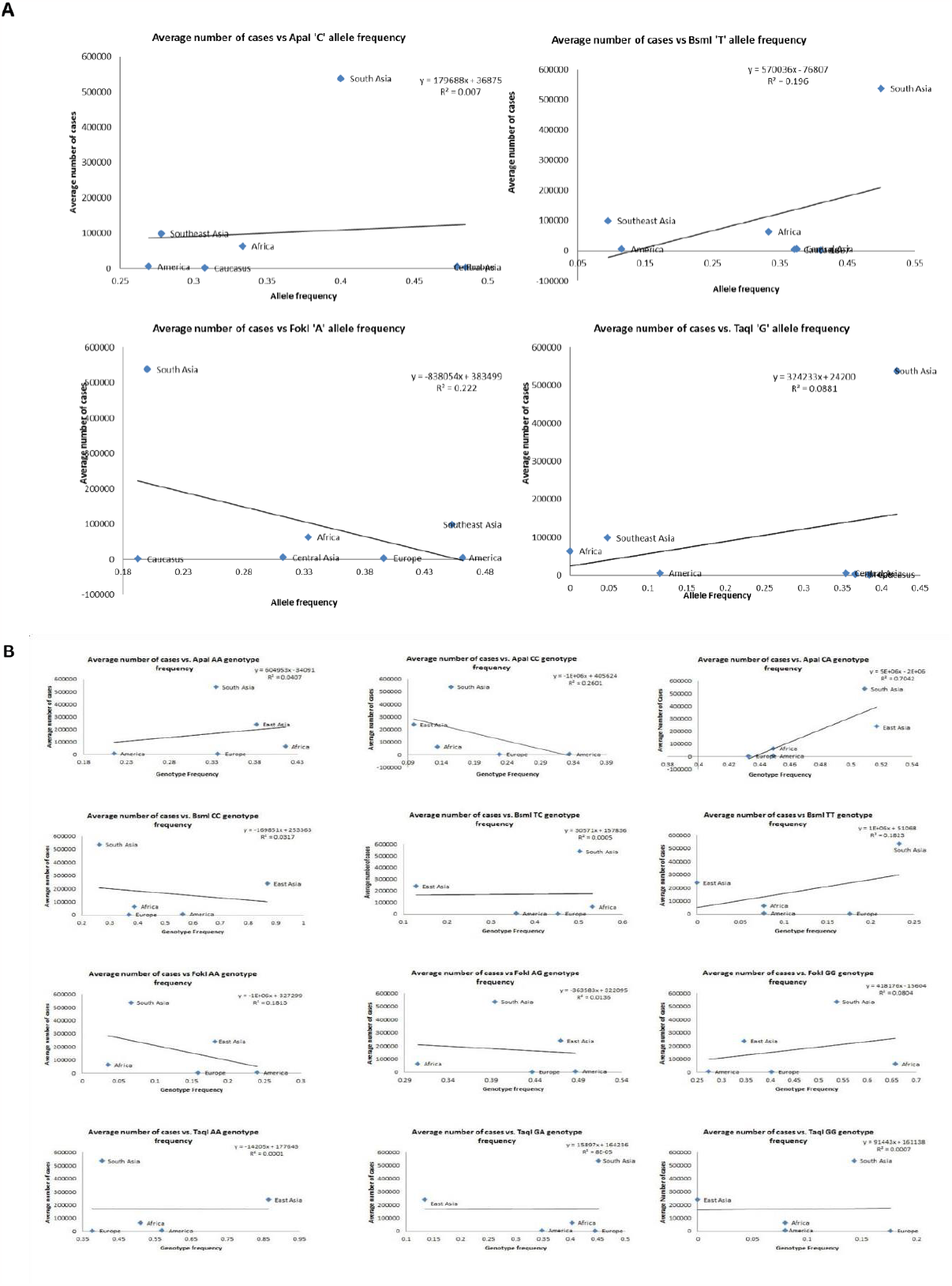
Graph showing correlation between prospective causative allele of the SNPs with the average number of cases. (b) Graph showing correlation between genotype frequencies of each SNPs with average number of cases in global populations.

## Discussion

Vitamin D is important not only for calcium homeostasis but also as an immune modulator as has been established by some previous studies. Vitamin D deficiency is regarded as a silent epidemic in most second and third-world countries around the globe and most patients with tuberculosis also present a low serum level of Vitamin D. As Vitamin D exerts its function through its receptor, i.e Vitamin D receptor (VDR), on the cell’s nucleus, it was hypothesised that the major polymorphisms of the VDR will alter the effectiveness of Vitamin D. There have been studies on the polymorphisms of VDR before especially the ones present on the 3’ untranslated region but no conclusive results were obtained. This could be because of insufficient sample size or differences in ethnicities.^[8,14,15,22,23]^

In our study, we explored that the four SNPs varied greatly in global populations. Despite the high D’ values of linkage disequilibrium between the SNPs, the r^2^ values were relatively less due to differences in allele frequencies of the SNPs within the same population group. From all the analysis we can conclude that the VDR polymorphisms are more variable in South Asians than in any other population. In other words, South Asians lack haplotypes where these SNPs are included as seen both in linkage disequilibrium studies and network analysis. Also, the pairwise differences within the population were highest in South Asians for all the SNPs. We also know that South Asians bear a high burden of the disease with India producing the maximum number of cases worldwide. The correlation graphs showed no association between the average number of cases and the causative alleles of the four polymorphisms. So it can be concluded that any SNP alone is at least not responsible for the manifestation of the disease. Similarly, none of the correlation coefficients between genotype frequency and the average number of cases reached a significant p-value.

## Supporting information

Linkage Disequilibrium 1000 genome

## Data availability statement

All datasets generated for this study are included in the article/Supplementary Material.

## Author contributions

GC and DD conceived and designed this study, analyzed the data and wrote the manuscript. All authors contributed to the article and approved the submitted version.

---

**Table 2:** Table showing correlation coefficients between average number of cases in global populations and prospective causative allele frequencies of the four VDR polymorphisms.

| Populations | Average number of cases | rs7975232 (APAI) C/A | rs731236 (TAQI) G/A | rs1544410 (BSMI) T/C | rs2228570 (FOKI) A/G |
| --- | --- | --- | --- | --- | --- |
| Africa (1KG) | 63421.29 | 0.3333 | 0 | 0.3333 | 0.3333 |
| America (1KG) | 5508.371429 | 0.2692 | 0.1154 | 0.1154 | 0.4615 |
| Caucasus(1KG) | 1667 | 0.3077 | 0.3846 | 0.4103 | 0.1923 |
| Central_Asia(1KG) | 5801.333333 | 0.4792 | 0.3542 | 0.375 | 0.3125 |
| Europe(1KG) | 3671.741935 | 0.4851 | 0.3663 | 0.3713 | 0.396 |
| South Asia(1KG) | 537746 | 0.4 | 0.42 | 0.5 | 0.2 |
| Southeast Asia(1KG) | 98924.77778 | 0.2778 | 0.04762 | 0.09524 | 0.4524 |
| Correlation Coefficient |  | 0.083637786 | 0.296782348 | 0.442733165 | -0.471187394 |
| R2 |  | 0.006995 | 0.3356 | 0.196 | 0.222 |
| P-value (two tailed) |  | 0.8585 | 0.1729 | 0.3199 | 0.2858 |
| P-value summary |  | ns | ns | ns | ns |

**Table 4:** Table showing the correlation coefficients between average numbers of cases in each populations along with the genotype frequencies of the four polymorphisms.

| Populations | Average Number of Cases (WHO, 2020) | ApaI |  |  | BsmI |  |  | FokI |  |  | TaqI |  |  |
| --- | --- | --- | --- | --- | --- | --- | --- | --- | --- | --- | --- | --- | --- |
|  |  | AA | CA | CC | CC | TC | TT | AA | AG | GG | AA | GA | GG |
| Africa | 63421.29 | 0.416 | 0.449 | 0.134 | 0.39 | 0.532 | 0.077 | 0.036 | 0.305 | 0.658 | 0.51 | 0.403 | 0.08 |
| America | 5508.371 | 0.216 | 0.449 | 0.334 | 0.564 | 0.357 | 0.077 | 0.24 | 0.487 | 0.273 | 0.57 | 0.348 | 0.08 |
| East Asia | 239146.3 | 0.382 | 0.517 | 0.099 | 0.871 | 0.128 | 0 | 0.182 | 0.47 | 0.347 | 0.865 | 0.134 | 0 |
| Europe | 3671.742 | 0.337 | 0.433 | 0.228 | 0.369 | 0.453 | 0.176 | 0.159 | 0.437 | 0.403 | 0.377 | 0.445 | 0.176 |
| South Asia | 537746 | 0.335 | 0.509 | 0.155 | 0.263 | 0.503 | 0.233 | 0.067 | 0.394 | 0.537 | 0.404 | 0.451 | 0.143 |
| Correlation Coefficient (Average number of cases with genotype) |  | 0.2017 | 0.8392 | -0.5100 | -0.1782 | 0.0219 | 0.4269 | -0.4257 | -0.1165 | 0.2835 | -0.0122 | 0.0091 | 0.0272 |
| R 2 |  | 0.04068 | 0.7042 | 0.2601 | 0.03175 | 0.000483 | 0.1823 | 0.1813 | 0.01358 | 0.08041 | 0.000149 | 0.000084 | 0.000744 |
| P-value |  | 0.7449 | 0.0755 | 0.3801 | 0.7743 | 0.972 | 0.4734 | 0.4748 | 0.852 | 0.6438 | 0.9845 | 0.9883 | 0.9653 |
| P value summary |  | ns | ns | ns | ns | ns | ns | ns | ns | ns | ns | ns | ns |

