## Supplementary figures and images for "Reassessing the association of VDR and its polymorphisms with tuberculosis in global populations"

### Supplementary Fig 1

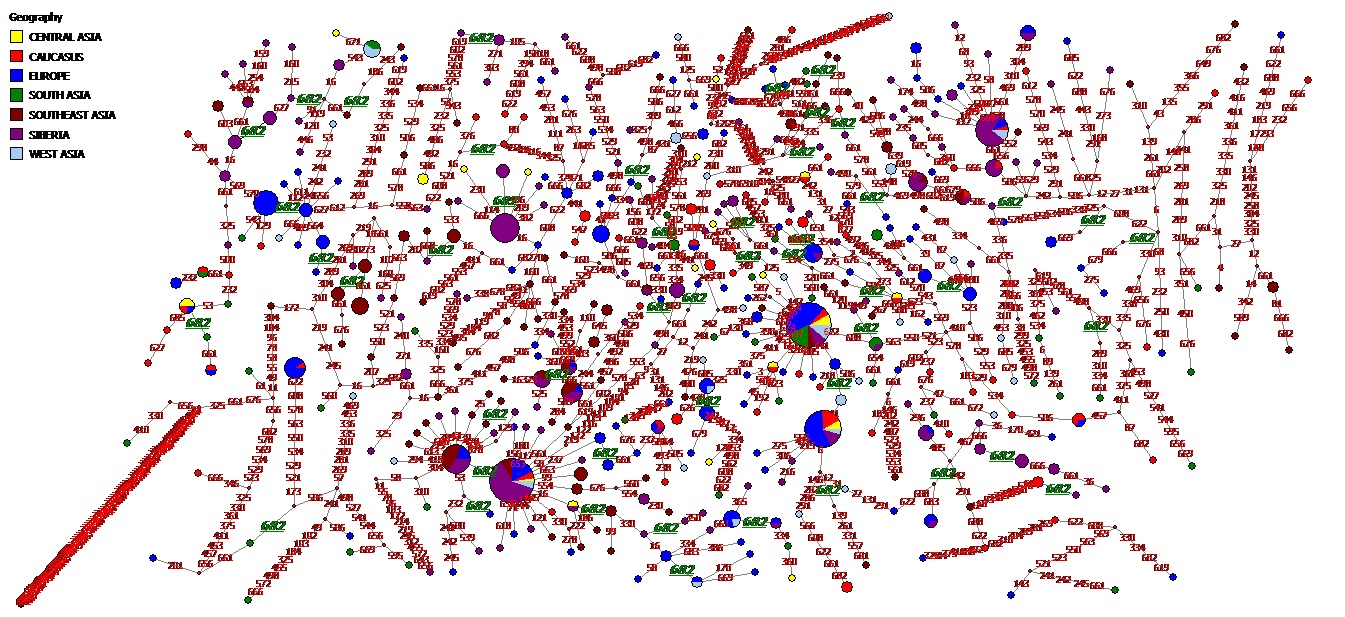

### Supplementary fig 2

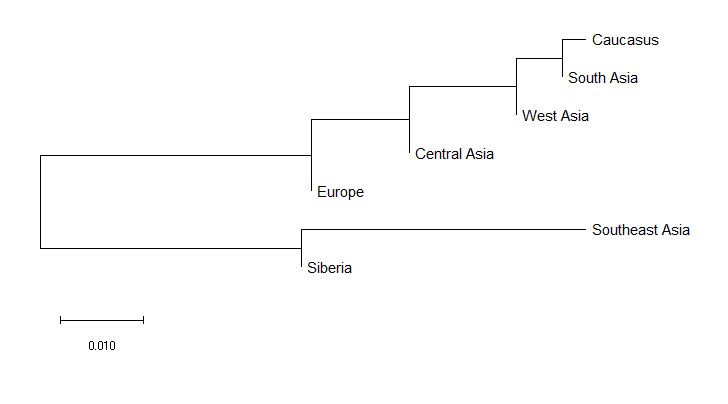
