## Supplementary Figure Legends for "Reassessing the association of VDR and its polymorphisms with tuberculosis in global populations"

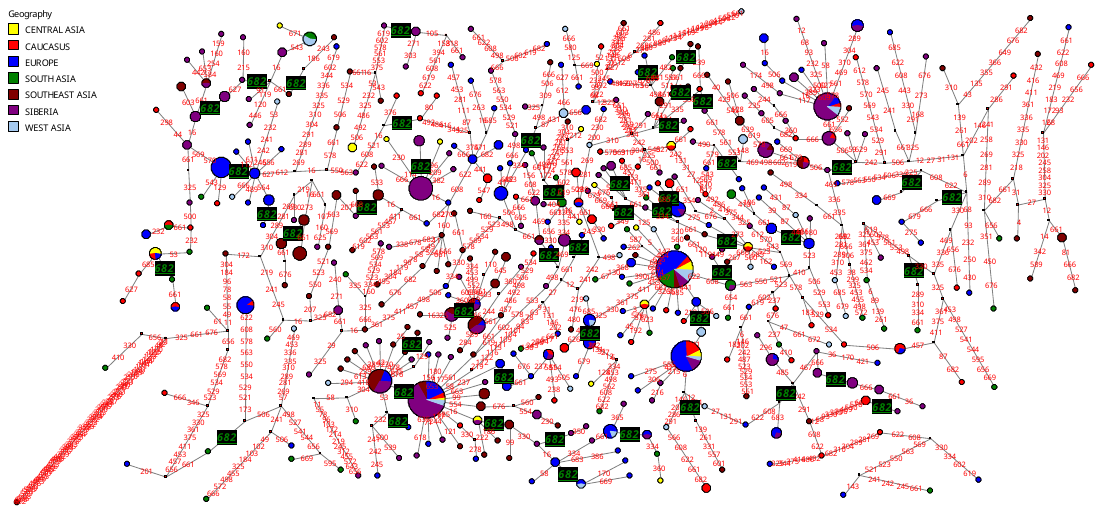

Supplementary Fig 1.: Median Joining Network of 30bp VDR sequence including rs2228570 (FokI). Here all the major haplotypes are seen as was observed in the main manuscript. The position of the hypervariable variant is highlighted here. It does not participate in any of the major haplotypes rather is involved in the formation of singletons giving rise to more diversity.

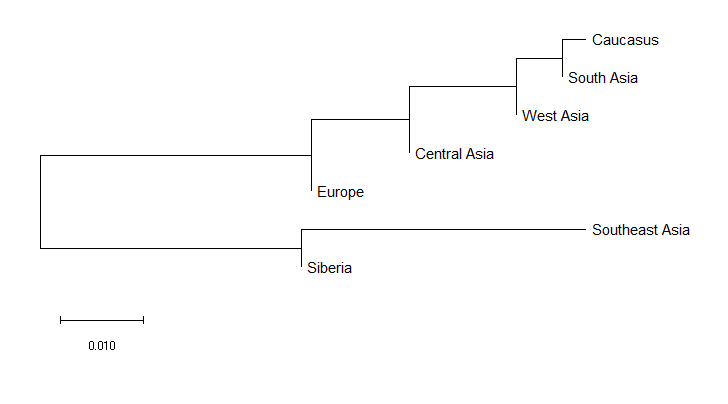

Supplementary Fig 2.: Neighbour-joining phylogenetic tree showing genetic relatedness between South Asia and West Eurasian population on the basis of the four VDR variants.
